# An antigenic diversification threshold for falciparum malaria transmission at high endemicity

**DOI:** 10.1101/2020.01.01.892406

**Authors:** Qixin He, Mercedes Pascual

## Abstract

In malaria and several other important infectious diseases, high prevalence occurs concomitantly with incomplete immunity. This apparent paradox poses major challenges to malaria elimination in highly endemic regions, where asymptomatic *Plasmodium falciparum* infections are present across all age classes creating a large reservoir that maintains transmission. This reservoir is in turn enabled by extreme antigenic diversity of the parasite and turnover of new variants. We present here the concept of a threshold in local pathogen diversification that defines a sharp transition in transmission intensity below which new antigen-encoding genes generated by either recombination or migration cannot establish. Transmission still occurs below this threshold, but diversity of these genes can neither accumulate nor recover from interventions that further reduce it. An analytical expectation for this threshold is derived and compared to numerical results from a stochastic individual-based model of malaria transmission that incorporates the major antigen-encoding multigene family known as *var*. This threshold corresponds to an “innovation” number we call *R*_*div*_; it is different from, and complementary to, the one defined by the classic basic reproductive number of infectious diseases, *R*_0_, which does not easily apply under large and dynamic strain diversity. This new threshold concept can be exploited for effective malaria control and applied more broadly to other pathogens with large multilocus antigenic diversity.

**Author summary:** The vast diversity of the falciparum malaria parasite as seen by the immune system of hosts in high transmission regions, underlies both high prevalence of asymptomatic infections and partial protection to re-infection despite previous exposure. This large antigenic diversity of the parasite challenges control and elimination efforts. We propose a threshold quantity for antigenic innovation, we call *R*_*div*_, measuring the potential of transmission to accumulate new antigenic variants over time. When *R*_*div*_ is pushed below one by reduced transmission intensity, new genes encoding this variation can no longer accumulate, resulting in a lower number of strains and facilitating further intervention. This innovation number can be applied to other infectious diseases with fast turnover of antigens, where large standing diversity similarly opposes successful intervention.

## Introduction

High transmission endemic areas present a significant challenge to the control and elimination of falciparum malaria due to the large reservoir of chronic asymptomatic infections sustaining transmission. Today the global burden of *Plasmodium falciparum* is concentrated in these high transmission endemic areas within fifteen countries, mainly in sub-Saharan Africa (WHO 2017). A similar reservoir is found in other vector-borne diseases that exhibit a high prevalence of infection with no clinical symptoms in domestic and wildlife hosts [1–3]. Large reservoirs of chronic asymptomatic infection arise not just from high transmission rates per se, but also from accompanying nonsterile specific immunity to pathogens with extreme antigenic variation encoded by multigene families [4, 5]. The accumulation and turnover of new antigenic variants which underlies such nonsterile immunity constitute a major impediment to intervention efforts for falciparum malaria [6].

One important multigene family in the malaria parasite *P. falciparum* is known as *var*. It encodes for the major antigen of the blood stage of infection, the protein PfEMP1 exported to the surface of infected blood cells upon expression. Anti-PfEMP1 immunity is crucial to prevent severe disease and clear infection [7]. Each parasite carries 40 to 60 var gene copies across its chromosomes. This variation enables immune evasion and associated long infection, as hosts have typically not seen many of the variant surface antigens of an infecting parasite. Under high transmission rates, local parasite populations exhibit a large pool of gene variants reaching the tens of thousands [8, 9]. This vast genetic diversity is generated primarily by ectopic recombination. Laboratory experiments have shown that a näive infection can generate about sixty new recombinants per year [10, 11], although this rate has not yet been demonstrated in nature. In addition, parasites share locally only a few common *var* genes across strains [9, 12, 13] and seasons [6]. Spatial diversity in *var* genes has also been documented [8, 9, 12], indicating that migration from surrounding areas also contributes to new diversity and to the immunological challenge.

A well-known and important quantity in epidemiology is the basic reproductive number *R*_0_, which establishes a threshold separating fundamentally different population dynamics in infectious diseases. The high strain diversity of *P. falciparum* and other pathogens with multigene and multilocus encoding of antigens [14–17] challenges the application of *R*_0_. Whereas its application to pathogens with either no antigenic variation or a low number of genetically well-defined strains is appropriate, its estimation and even definition becomes difficult when antigenic variation is large and dynamic under extensive recombination and no stable strains. Under these conditions, it is informative to consider the accumulation and turnover rate of new genes encoding for antigens, as their diversity can influence transmission characteristics and responses to control.

We present here a complementary number that defines a threshold for parasite antigenic diversification. Below this threshold, the accumulation of new antigen-encoding genes no longer occurs, even though they are consistently produced. We introduce the concept for infectious agents in general, derive an analytical expectation for the rate of generation of “successful” new genes for the *var* system in *P. falciparum*, and demonstrate the existence of the predicted analytical threshold in numerical simulations of a stochastic agent-based model that incorporates *var* genes and the acquisition of immunity by individual hosts. We then investigate the epidemiological and evolutionary factors that influence this diversification rate analytically. We show that this rate for the accumulation of genetic novelty, we call *R*_*div*_, maps onto transmission intensity, separating at a threshold a regime in which new genes are able to accumulate from one in which they are unable to do so despite transmission still occurring (i.e., *R*_0_ remaining above one). We discuss implications for malaria control and elimination, future directions to estimate and monitor distance to this quantity in high transmission endemic regions, and its applicability to other infectious diseases.

## Results

### Theoretical considerations

Consider the genes of a parasite whose mutation generates new antigenic variation during transmission and infection. Due to immune selection, common antigenic variants would segregate around a frequency of 1*/G* given an antigenic pool size *G*. We define the establishment of a new antigenic variant as its propagation from a few copies to the expected common variant frequency. Parasite populations should accumulate these new gene variants when they are produced at a sufficient rate for their lifespans to overlap with each other (Fig. 1). Novelty per se guarantees neither the establishment nor the persistence of the genes. Even under high absolute fitness, new variants need to survive initial drift to establish in the parasite population [18]. Their accumulation further requires that the rate at which they are generated, *G*_*new*_, be on average larger than that of their loss, given by the inverse of their lifespan, *T*_*new*_. In other words, at least one beneficial gene needs to be produced and become established in the population during the typical lifespan of a previously generated new gene. We denote by *R*_*div*_ the expected number of new genes established during the average lifespan of a new gene. This innovation number should be greater than 1 for new variants to accumulate, namely

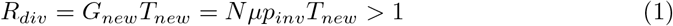

where *N* denotes the population size of the parasite (i.e., the total number of infections), *µ*, the mutation rate of the genes, and *p*_*inv*_, the invasion probability of a low-frequency variant. Note that *N* is different from the number of infected hosts because there could be multiple infections within the same host. Importantly, the average lifespan of a new gene, *T*_*new*_, depends on its selective advantage under frequency-dependent competition [13, 19]: if fewer hosts have acquired immunity towards its product, then its expected lifespan would be longer. New variants are generated through either mutation or ectopic recombination (Methods), and therefore *µ* generically refers here to the rate of novelty generation regardless of specific mechanism. Note that *R*_*div*_ is defined as an instantaneous measure of the establishment rate of new genes. As such, it does not concern the death rate of pre-existing common genes and is not a turnover rate. As we discuss later, the lifetime of pre-existing common genes under frequency-dependent selection is much longer than the epidemiological time scales relevant to the invasion of new, rare ones.

**Fig 1.**
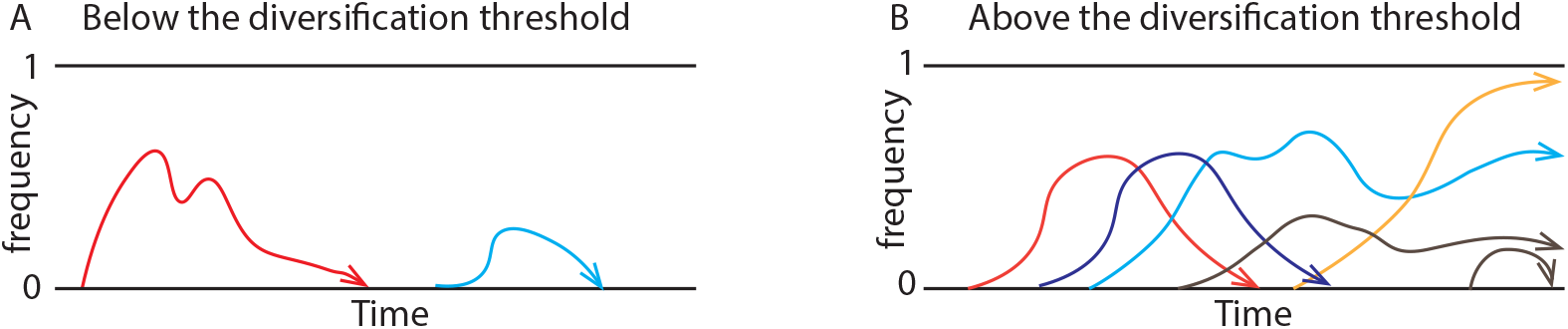
Schematic illustration of two possible scenarios. Below the diversification threshold, new genes are produced but die before any others arise (**A**). Above this threshold, they are produced within the lifespan of other new genes, causing their accumulation (**B**). Constant Red Queen dynamics between parasite antigens and the adaptive immune system under high transmission intensity, result in high diversity of antigen-encoding genes.

Equation (1) establishes an expectation for the existence of a threshold, whose expression we proceed to further develop next, to be able to verify it computationally. We first consider the probability *p*_*inv*_ that a new gene survives its initial low frequency and invades. Based on birth-death processes in the Moran model with selection [20], the invasion probability of a low-frequency variant is largely determined by its relative fitness advantage over other variants. The fitness of different variants in a transmission model is essentially given by their effective reproductive number *R*_*eff*_ (i.e., the number of infections they produce via transmission events during their lifetime). Thus, 

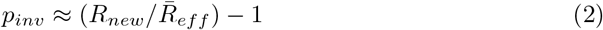

Equation (2) holds in general for any infectious disease that generates new antigens. To proceed further, we considered the specifics of *Plasmodium falciparum* and its multicopy *var* genes (typically about 40-60 per genome), whose expression is sequential during the blood stage of infection [21]. *R*_*eff*_ for a given *var* gene is the product of the epidemiological transmission rate of the disease (*β*) and the typical infection duration (*τ*) of parasites that carry the given gene (Methods). If we consider that genes are equivalent in transmissibility (i.e., their products are functionally equivalent in their ability to bind host receptors), an assumption we later relax in numerical results, fitness differences between variants are only determined by the duration of infection these genes can typically sustain.

Because only those genes whose product the host has not yet built immunity towards are expressed, the average duration of infection will equal the number of genes per genome (i.e., repertoire size) times the average proportion of susceptible (non-immune) hosts per gene, 

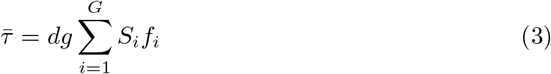

where *d* is the duration of infection for a given gene in a naive host, *g* is the repertoire size, *G* is the number of different genes in the parasite population (i.e., gene pool size) and *f*_*i*_ is the population frequency of a given gene. We rewrote equation (2) using (3) (Methods), to obtain 

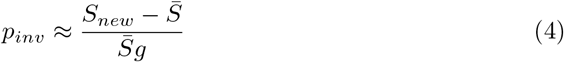

where the mean number of susceptible hosts for an average established gene is given by 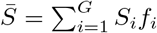. This expression for the invasion probability shows that a new gene is likely to invade when it affords a wider host niche than that of established genes (by encoding for epitopes that are new given the immunity of the host population). In other words, the available number of hosts for its expression should be higher on average than that for existing genes. In addition, the invasion probability of a single gene decreases with increasing repertoire size *g*, as the importance of a single gene also decreases.

In order to understand how epidemiological and genetic factors influence *R*_*div*_, we examined with a simple theoretical model the equilibrium values of 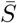 and *N*, which enter prominently in the expression for *G*_*new*_ (Methods). At equilibrium, increasing contact rates (*β*) via mosquito bites, result in higher parasite population sizes (*N*) and a lower average number of susceptible hosts (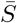, S1 Fig). More specifically, 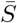 is mostly determined by *β* and *g*, whereas parasite population size strongly scales with the diversity ratio (i.e., gene pool size *G* over repertoire size, *G/g*). A lower 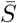 favors invasion of a new variant (i.e., increases *p*_*inv*_), and a higher parasite population size (*N*) and a genome with a higher number of unique genes (*g*) generate new variants faster. Note that theoretical predictions underestimate 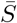 because they neglect the higher-level of organization of the genes into different genomes, also under immune selection [13]. This observation indicates that strain structure under selection can significantly reduce the percentage of genes that a host is immune to, especially under high competition and high diversity (see *g* = 60, and *G/g* = 100 in S1B Fig).

An explicit expression for *T*_*new*_ cannot be obtained analytically because this quantity continuously changes as new genes enter the system and influence the nonlinear dynamics of *N, G*, and 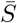. To gain nevertheless an understanding for how these variables affect *T*_*new*_, we approximated the average lifespan of a new gene on the basis of an adapted diffusion equation [20] (SI). The diffusion equation for 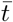 requires consideration of how the frequency of a new gene *x*(*t*) varies in time (SI). When applied to our model, the resulting analytical approximation for 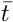 (Eq. S8-11) shows that the expected lifespan of a new gene grows faster than exponential with decreasing 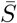, and surpasses the average time to fixation of a neutral gene (2*N*) when 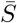 is below a given value (*∼* 40%) (Fig. 2).

**Fig 2.**
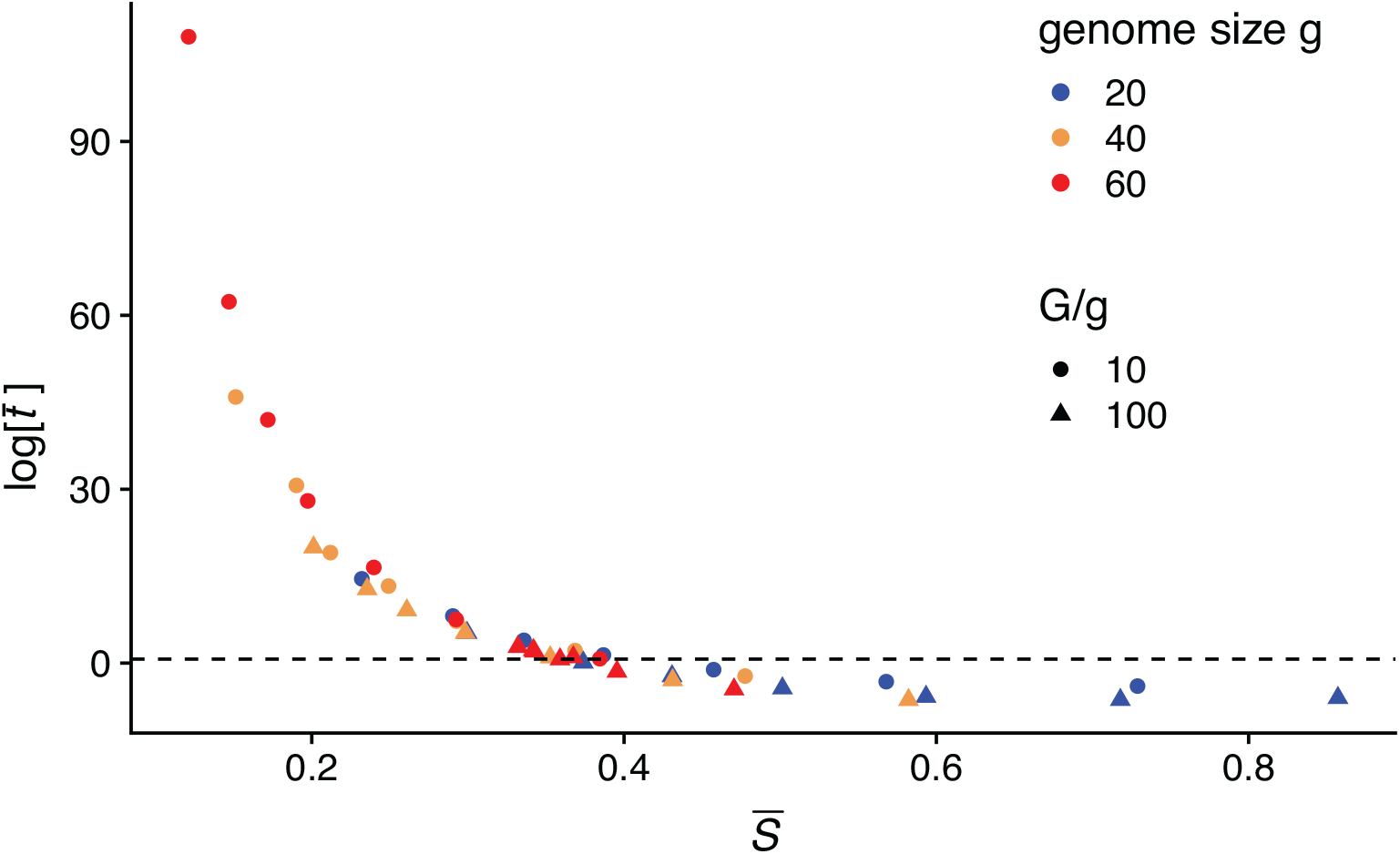
Theoretical expectation of the average lifespan of a new gene 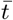. The analytical expression shows that 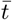, measured in units of 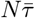 increases faster than exponential as the average number of available hosts 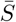 decreases. The dashed line represents the time to fixation of a neutral gene, which means that under small 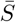, once established, the gene can be maintained in the population for much longer than the typical epidemiological timescale (or for much longer than the simulation period of 200 years in our model). The average lifespan *T*_*new*_ obtained from the computational model will always be considerably smaller than the theoretical expectation 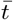 derived under the assumption that other factors remain constant, in particular the average number of hosts 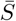 that are susceptible to the invading gene. The general trend of increased persistence with lower 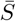 will hold however for the full numerical system and for finite time windows.

Taken together, the theoretical results show that when transmission rate is low, 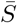 is large and new genes do not have a significant advantage over older ones (S1 Fig). New genes experience a small invasion probability, and even when they invade, they experience strong drift, functioning as effectively neutral (Fig. 2). As transmission intensity increases, the selective advantage of new genes also increases as 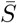 decreases. Once 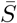 is below a given value (0.4 in our simulations), new genes are most likely to be maintained in the population indefinitely. Concomitantly, the increase in gene diversity results in higher parasite population sizes *N* (S1A Fig). The system thus enters a regime of positive feedback for new variants, as elevated diversity boosts *N* and therefore also, *G*_*new*_, before reaching equilibrium.

### Computational model and threshold implications for epidemiology

We can now evaluate the existence of the threshold behavior indicated by the above analytical argument. To this end, we computed *R*_*div*_ from a stochastic agent-based model of malaria transmission [13, 22] that tracks *var* evolution and immunity and is described in Materials and Methods [13]. A number of extensions were also considered here to address the generality of the argument, including distinct major *var* gene groups with associated differential fitness and constrained recombination (Methods). Note that all the simulations are endemic, with *R*_0_ larger than 1. Although *R*_*div*_ is defined as an instantaneous measure, we estimated its mean value from the simulations over a given period of time because of the non-linear dynamics of the system, which modify components of the number as new genes continuously invade. We then examined how the accumulation of new variants over this period of time varies as a function of *R*_*div*_. We calculated *G*_*new*_ according to Eq. (2) and (4) by obtaining 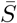, *N*, and *µ* directly from the simulations, and *T*_*new*_ from the average lifespan of all the new variants that are produced during this time period. The mean *T*_*new*_ evaluated numerically will always be shorter than that predicted from the diffusion approximation 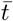, especially as 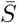 decreases and persistence times rapidly increase (Fig. 2). This numerical value is smaller because the stochastic simulations can only track lifespan within a finite time period (which places an upper bound on its value), and because the assumption of constant 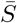 does not apply. Nevertheless, the theoretical trend of increasing *T*_*new*_ with decreasing 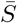 and therefore, higher transmission intensity, does apply to the numerical system.

Results showed that the transmission system naturally falls into two regimes separated by a threshold at *R*_*div*_ = 1 (Fig. 3A). Below the transition, new antigenic variants are generated but do not accumulate or persist (Fig. 4A), whereas above it, they are able to accumulate and experience a continuous turnover rate (see shifting shades of colors in Fig. 4B). The transition between these regimes occurs around the proposed boundary where the rate of generation of genes surpasses the average lifespan of new antigenic variants (*R*_*div*_ = 1). This threshold is robust to differences in specific assumptions about the transmission and genetic systems (including processes of within-host dynamics, functional differences between genes, values of the recombination and biting rates), as each point in Fig. 3A represents a simulation with different model assumptions and parameter combinations (Methods; S1-2 Table).

**Fig 3.**
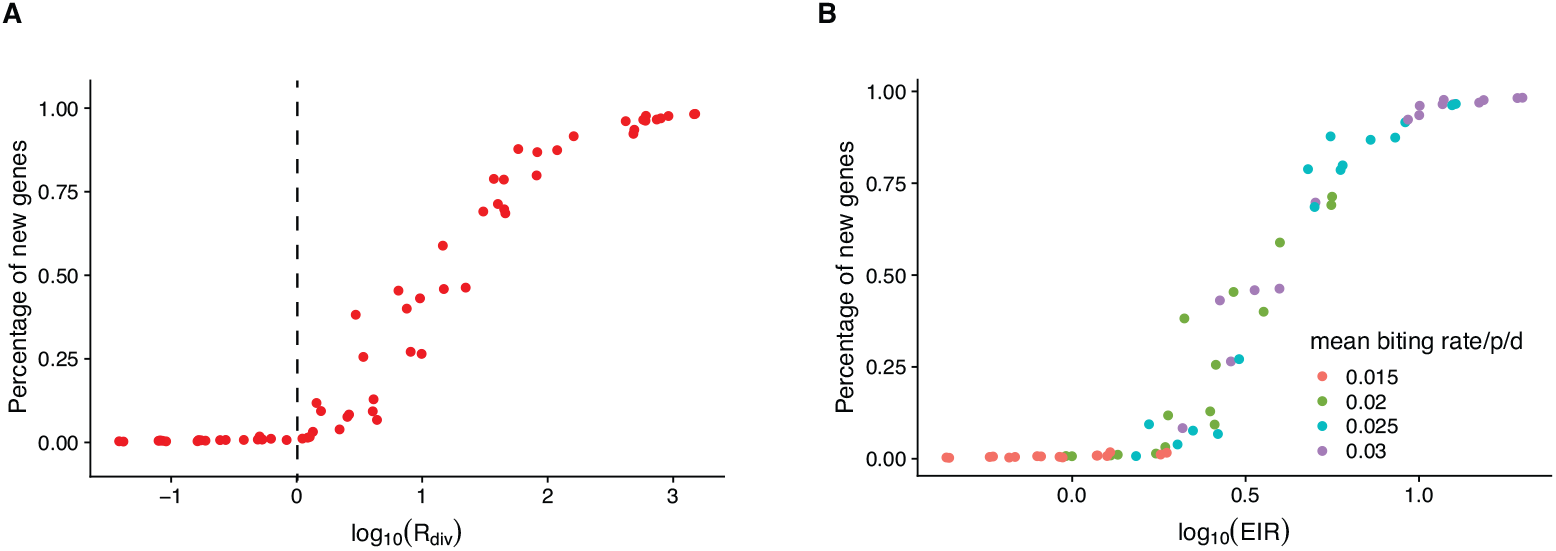
Numerical simulations reveal a transition between two regimes of antigenic diversity accumulation. (**A**) The percentage of new genes in the local parasite population at the end of a given simulation period (200 years) remains negligible when the reproductive number *R*_*div*_ for antigen-encoding new genes is lower than one. By contrast, this percentage increases rapidly above this threshold. Because the time interval over which we computed *R*_*div*_ = *G*_*new*_*T*_*new*_ concerns long transients, we evaluated the rate of generation of new genes *G*_*new*_ as a mean over this interval (by averaging the values of *N* and 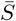 every 180-day interval), and the lifespan *T*_*new*_, as an average for all the new genes that invaded during this time (with this interval placing an upper bound on individual lifespans). (**A, B**) Each point represents a simulation with different combinations of parameters and assumptions (including variation in rules of within-host dynamics, in strength of the trade-off between transmissibility and duration of infection, and in values and seasonality of the transmission rates, Table S1-2). (**B**) The percentage of new genes also exhibits the threshold behavior with the entomological inoculation rate (EIR, the number of infectious bites per person per year).

**Fig 4.**
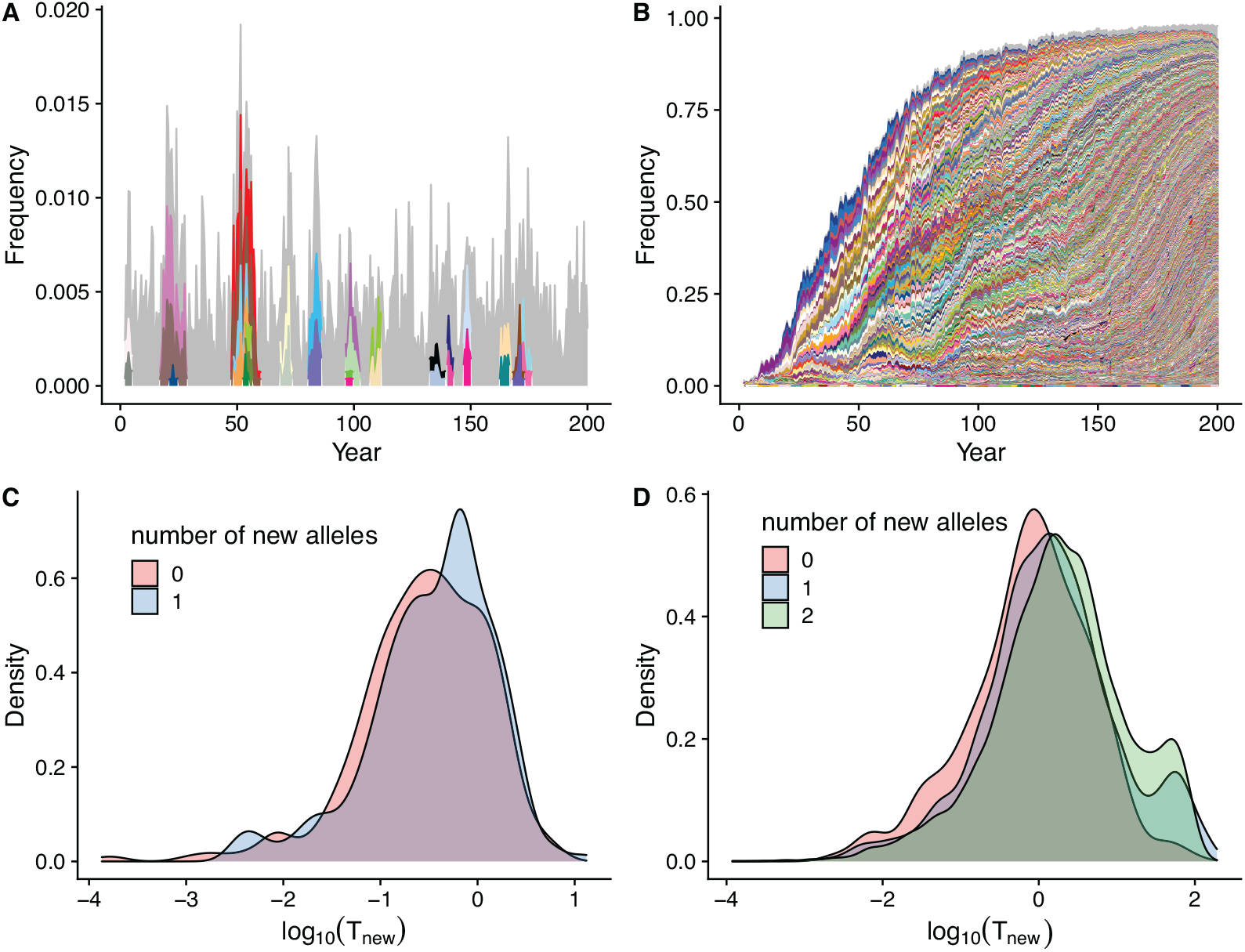
Patterns of new gene establishment in simulations below and above the *R*_*div*_ threshold of 1. (**A**) New genes do not accumulate below the transmission threshold where they essentially follow neutral dynamics. In contrast, they do accumulate at a constant rate under frequency-dependent selection above this threshold (**B**). Each color in the stacked area plots refers to a new gene in the population. New genes that account for less than five infections over the entire period are combined and represented in grey. (**C**) The distribution of lifespan of new genes generated in the simulation (measured in *log*(*years*)) is shorter below the threshold than above it (**D**). In addition, new genes with a greater number of new epitopes live longer above the threshold, whereas below the threshold, they experience similar lifespans (**C, D**).

Importantly, we found that the quantity *R*_*div*_ scales monotonically with the intensity of transmission measured here as the entomological inoculation rate (or *EIR*, the number of infectious bites per person per year) (S2 Fig), a practical empirical measure from field epidemiology. The association with *R*_*div*_ should hold more generally with any other measure of transmission intensity. This association implies that the transition between regimes also occurs as a function of transmission intensity (Fig. 3B), and therefore, that the malaria system can be pushed below threshold by changing this control variable. It follows that the percentage of accumulated new genes also exhibits the threshold behavior with transmission intensity, as measured here by EIR (See S2 Fig caption for more details).

The transition examined so far represents the behavior of the system for different values of *R*_*div*_ or transmission intensity. Its existence should influence the temporal response of the malaria system to intervention events that reduce transmission at a given point in time. In particular, interventions that take the transmission system below threshold should lead to distinct responses than those that fail to do so. This is illustrated in Fig.5 A, where we numerically introduced a transient reduction of the biting rate to lower levels (respectively 30 and 50% of its original value). New genes cease to accumulate only when *R*_*div*_ goes below threshold, whereas they continue to invade and accumulate following a temporary decrease otherwise (Fig. 5A). We further investigated how the accumulation of new genes influences the outcome of intervention with respect to malaria incidence. When intervention reduces transmission rates, both prevalence and MOI decrease, whereas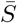 increases, as expected (Fig.5B, C). Importantly, *R*_*div*_ predicts whether prevalence and MOI would be able to recover through the accumulation of antigenic variation. When *R*_*div*_ remains larger than 1, the system is able to partially recover both its prevalence and MOI in a relatively short period of time depending on the actual *R*_*div*_ values. In contrast, when *R*_*div*_ is taken across threshold to values smaller than 1, the system stays at the low prevalence determined by control levels and does not rebound (Fig. 5 B, C). It is not possible to determine the trajectories of these epidemiological quantities from their values at intervention, or from monitoring the incidence rates themselves, in the sense that there is no given, knowable threshold in these quantities that would indicate the discrete transition.

**Fig 5.**
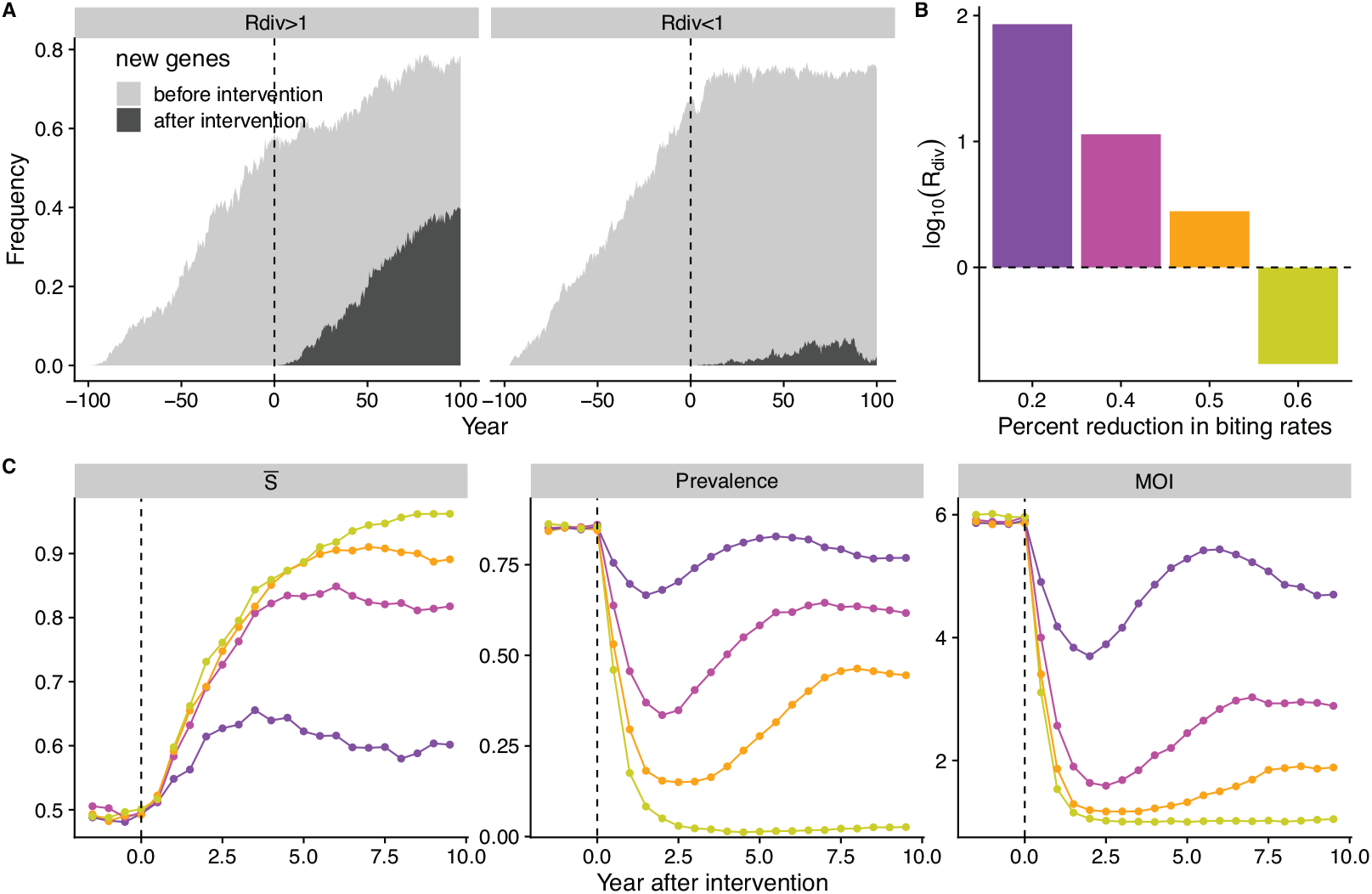
*R*_*div*_ predicts the response of incidence to reductions of the transmission rate. (**A**) Interventions that push the system below the threshold are effective at stopping the accumulation of new genes, whereas those that do not, result in the rebound and rebuilding of diversity (**A**). Light and dark grey colors indicate genes that originate respectively before and after the intervention. Reductions of transmission rates would result in decreased *R*_*div*_ values and increased 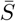 (**C**) to different levels. (**C**) When *R*_*div*_ remains above 1, prevalence and Multiplicity of Infection (MOI) rebound relatively fast, whereas they do not recover when *R*_*div*_ drops below 1. Changes in these epidemiological quantities, prevalence and MOI, cannot per se indicate this transition, as they change continuously with different levels of intervention and do not exhibit a threshold.

In summary, by evaluating whether *G*_*new*_ *<* 1*/T*_*new*_ (or equivalently, *R*_*div*_ *<* 1), one can predict whether the system has a relatively stable antigen-encoding gene pool, or whether alternatively, new variants continuously enter into it. We have shown that whether new genes can successfully establish in the population is most tightly linked with the average proportion of susceptible hosts (or the niche) available for existing genes 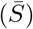, a measure of the population-level immunity. In addition, this measure provides a valuable assessment tool for evaluating the potential success of interventions by predicting whether incidence rates would rebound.

## Discussion

The concept of *R*_*div*_ arises from the interplay of immune memory and antigenic variation at the population level, as a result of frequency-dependent selection. As such, it differs from the antigenic diversity threshold previously proposed for the HIV virus and its transition to AIDS, arising from the race between viral replication and immune responses at the within-host level [23]. The concept itself and the associated transition regime described here should apply more generally to other infectious diseases with antigen-encoding multigene families, such as *vsg* genes in *Trypanosoma brucei* and *msg* genes in *Pneumocystis carinii* [4]. Because the basic concept is independent of specific consideration of multigene families and their properties, it should also be adaptable to other pathogens in which large standing antigenic diversity at the population level results from multilocus genetic variation [17].

By contrast, in pathogens with sufficiently well-defined strains, the characteristics of their population dynamics and population genetics would place them below the diversification threshold defined by *R*_*div*_ equal one. *R*_*div*_ does not provide additional information in these systems and estimates of the force of infection through tracking the transmission of new clones [24] would be sufficient. For example, genetic variation in measles is largely neutral antigenically and the effective mutation rates generating new antigens are slow [25]. In seasonal influenza, bottlenecks in transmission constrain the emergence of novelty [26], and so do mutations with largely deleterious effects [27].

The innovation number *R*_*div*_ measures how many rare genes become frequent and established in the average lifetime of newly-generated genes. In contrast to *R*_0_ *<* 1 which predicts a decrease in parasite population size, *R*_*div*_ *<* 1 implies a stasis in overall antigenic diversity. This is so because *R*_*div*_ does not take into account the loss of established common genes, which occurs rarely given that they can persist under frequency-dependent selection for much longer times than neutral ones, even in a relatively small population [28, 29]. These persistence timescales are considerably longer than the epidemiological ones we are interested in here for the invasion and accumulation of novelty, and therefore our formulations have essentially assumed a separation of these timescales. Below threshold, the number of new genes would decrease if one were to track the fate of a given number of them introduced at low frequencies characteristic of invasions.

For *P. falciparum* and pathogens with extensive antigenic diversity at the population level, the proposed concept of a threshold behavior in the accumulation of antigen-encoding genes has practical implications for overcoming the resilience of highly endemic regions to intervention efforts. Although a decreasing trend in the diversity of strains and underlying genes with decreasing transmission intensity is well known and expected from both the biogeography and epidemiology of malaria, the actual form of this relationship and response to interventions are much less clear. Our results predict the existence of a sharp transition below which the disease system should effectively respond as a typical low transmission region, not just because of reduced transmission intensity but also because of much lower antigenic diversity no longer able to rebuild. Failure to push transmission intensity below this threshold would lead to a fast rebound in new antigenic variation and the recovery of prevalence and MOI. The crossing of the threshold would instead provide an indication that the system is now poised for further intervention with enhanced results. Thus, our new quantity can help evaluate an aspect of intervention that remains hidden on the basis of typical epidemiological quantities.

Control and even elimination efforts are indeed known to be most successful in biogeographical regions of low transmission, such as those at the edge of the distribution of the disease in Africa and in other continents [30]. Arresting the fast turnover of the local antigenic pool typical of high endemism would significantly repress disease burden and facilitate its further reduction. Concomitant control efforts at a regional level are critical to stem immigration, as migrant genomes would exhibit higher invasion probabilities than local ones, given their higher likelihood of encoding new antigens. Monitoring the turnover of *var* gene diversity through molecular epidemiology in response to control efforts should inform intervention evaluation in high transmission regions.

Although the importance of host immune selection in shaping the antigenic variation of *P. falciparum* and other pathogens is recognized [19, 31–33], mathematical and computational models typically evaluate intervention efficacy without explicit consideration of antigenic diversity (e.g. [34, 35]) and openness of the system to innovation [36]. Our results underscore the importance of these aspects.

Estimation of the diversification number, *R*_*div*_, would provide general guidelines for intervention evaluation where monitoring incidence rates alone would not suffice. Future work should consider how to obtain this number from an estimation of key parameters, including parasite population size, transmission rates, and gene pool size, based on combined data from molecular and field epidemiology. Parameterization of an agent-based stochastic transmission model that implements immune selection and recombination explicitly (e.g., [13, 37]) could be used, which represents a computational challenge (but see [38]). Estimating the viability of new recombinants will require bioinformatic analyses of population-level *var* sequence data for the *DBL*_*α*_ portion of the gene [39].

For simplicity, our analytical derivations treated each gene independently, even though *var* gene composition in parasite genomes of local populations in regions of high transmission has been shown to be non-randomly and non-neutrally structured, exhibiting low overlap as the result of immune selection [9, 13, 22, 40]. Hence, the fate of a viable new antigen-encoding gene depends on its genomic background, which ultimately determines the strength of competition among parasites. Comparisons of analytical expectations with numerical simulations revealed an influence of such population structure on the fate of new genes, and therefore, on components of *R*_*div*_. Future work should examine extensions of this work that account for this further complexity of immune selection operating at different levels of organization. In addition, our model does not consider cross-immunity or other types of generalized immunity, which have been proposed to prolong chronic infections of malaria [41, 42]. Because these forms of immunity do not directly influence whether antigenic diversity is at a selective advantage or not, we focused on variant specific immunity and on parameters that would influence the generation and maintenance of *var* encoded antigenic diversity. Other types of immune memory would influence strains as a whole beyond their *var* identity. Future work will examine further aspects of immunity.

In summary, explicit consideration of a threshold in antigenic diversification should enhance our understanding of transmission dynamics where large standing pathogen diversity represents a major challenge to control efforts.

## Materials and methods

### Analytical derivation of R_div_

We consider a population of hosts whose number is denoted by *N*_*host*_, receiving malaria infections from a diverse set of parasites, each composed of *g* genes from a constant gene pool of size *G*. One of the two main components of *R*_*div*_ is the rate at which new genes are generated, *G*_*new*_. Besides the mutation rate *µ* and the equilibrium parasite population size *N*, its expression requires the invasion probability *p*_*inv*_, we derive below.

#### Invasion probability of a new variant, *p*_*inv*_

From birth-death processes according to the Moran model with selection [20], the probability of establishment of a low frequency variant is determined by its fitness advantage relative to that of other genes, and by the parasite population size. That is, 

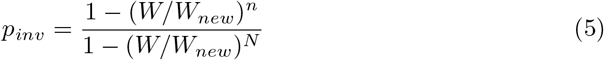

where *n* denotes the number of copies of new genes, and *W*, the fitness of a gene. In our case, *n* = 1 as new genes originate from a unique mutation or an ectopic recombination event. When *N >>* 1, the invasion probability *p*_*inv*_ is approximately its initial selective advantage relative to established gene variants, provided the selection coefficient remains the same. Since the fitness of each individual gene in a transmission model is essentially given by their effective reproductive number *R*_*eff*_ for existing genes and *R*_*new*_ for the new genes, we have 

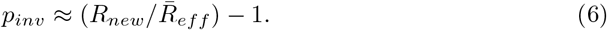

*R*_*eff*_ for a given *var* gene is in turn the product of the epidemiological contact rate of the disease (*β*) and the typical infection duration (*τ*) of parasites that carry the given gene, 

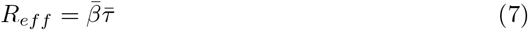

The contact rate *β* is equal to the product of the transmission rate (*b*) and the ‘transmissibility’ or infectivity of the given *var* gene (i.e., the functionality of the gene, *f*). Because we do not model vectors explicitly in the numerical stochastic model, the contact rate (*β*) refers to the rate at which a transmission event occurs, with a ‘donor’ host transmitting infection to a ‘recipient’ one (detailed description in section on “the modified *var* evolution model”).

Different groups of *var* genes may vary in their binding affinities to host receptors and therefore in their transmissibility. For simplicity, we consider that all genes exhibit the same transmissibility and therefore, the same absolute fitness, as we are most interested in estimating the fate of a new variant as a result of immune (frequency-dependent) selection. (We do explore later the effect of fitness differences numerically with the agent-based stochastic model, described in the section on “the modified *var* evolution model”).

With (9) for *R*_*eff*_, we can write 

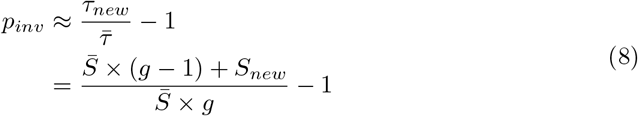

#### Numerical evaluation of *R*_*div*_

With equation (8) (or its equivalent (4)), we can now compute *G*_*new*_ = *Nµp*_*inv*_ in equation (1) from the output of our numerical simulations (described below under the *The modified var evolution model*). We also obtained from the simulations the other major component of *R*_*div*_, the mean lifetime of the genes in the system, *T*_*new*_, by directly tracking their fate individually. Although an analytical expression for *T*_*new*_ was not achievable for this nonlinear and stochastic transmission system, we considered gene lifetime under simplifying assumptions, as explained next.

#### Analytical derivation of 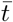

The simplifying assumptions are that the system has reached an equilibrium ((i.e., parasites get transmitted and die at the same rate), and that only the average lifetime of a gene varies, with all other variables remaining unchanged, including *N* and the average proportion of hosts 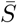 susceptible to an average gene. To differentiate gene lifetime under these conditions from *T*_*new*_ itself, we call it 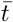. An expression for 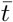 is derived by considering the frequency-dependent selection experienced by a new gene variant entering the system at equilibrium. We specifically approximate the dynamics of 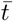 on the basis of an adapted diffusion equation [20] (SI).

## The modified *var* evolution model

We used an extended implementation of the agent-based model developed in [13]. Here, we first describe the specifics of the computational model, and then document the specific changes implemented in this study, including different transmission scenarios and rules of within-host dynamics. (Parameter combinations and specific rules are listed in Table S1-2).

The computational model is an individual-based, discrete-event, continuous-time stochastic system in which the infection and immune history of each host are tracked individually. Each infection object consists of a *var* repertoire, an infected host, the order of gene expression, and the timings of next events, including transition to expression of the next *var* gene, mutation/recombination, and clearance. Upon transition to a different *var* gene or clearance, the host gains specific immunity towards the epitopes in the gene. Global events include local transmission from biting events, new transmission from migrant *var* repertoires, and birth and death of hosts. In the numerical implementation of the simulation, all possible future events are stored in a single event queue along with their putative times, which may be fixed or drawn from a probability distribution. When an event occurs, it may trigger the addition or removal of future events on the queue, or changes of their rates, leading to a recalculation of their putative time. This implementation is adapted from the next-reaction method following [43], which optimizes the Gillespie first-reaction method [44] and allows for faster simulation times.

### Transmission and within-host dynamics

Local transmission events are sampled at the rate, *N*_*host*_*β*, in which a donor and a recipient host are sampled randomly from the host population. If the donor harbors parasites, then each parasite has a probability of being transmitted to the mosquito that is proportional to the functionality of the *var* gene that is currently under expression. *var* repertoires picked up by the mosquito will recombine with another genome to produce sporozoites. Specifically, if there are *n* parasite genomes, each genome has a probability 1*/n* of recombining with itself, producing the same offspring genome, and a probability 1 − 1*/n* of recombining with a different genome, producing recombinants. The total number of *var* repertoires passed onto the receiver is kept the same as that received from biting the infectious host. The life cycle of each *var* repertoire encompasses the liver stage, asexual blood stage, and the sexual stage. Parasites in a human host are infectious only at the asexual stage. Since we do not model mosquitoes explicitly, we implement a delay of 14 days between a transmission event and the repertoire becoming infectious, representing altogether oocysts development in the mosquito during the sexual stage and the initial liver stage in the receiver host. *Var* genes within a repertoire express sequentially in a random order, or according to their functional level from high to low, depending on the specific rule (Table S1), with a switching rate of 1*/d* (with *d* denoting the mean duration of expression). If the host has prior immune memory to the gene, the expression switches to the next gene instantaneously. When the expression switches to a new gene, immunity to the gene previously expressed is added to the host’s immune memory. The infection ends when the expression of the whole *var* repertoire is completed. Therefore, duration of infection, *τ*, is not varied a priori as a function of age but determined by the number of new epitopes to a given host. The immunity towards a certain epitope wanes at a rate *d* = 1/100 per day [45]. Mutation *µ* and ectopic recombination *r* occur randomly during the infectious stage (see below).

### *Var* repertoire structure

The repertoire of an individual parasite is a combination of *g* var genes. Each *var* gene, in turn, is a linear combination of two loci encoding epitopes that are connected linearly, and each epitope can be viewed as a multi-allele locus with *n* possible alleles. The initial conditions for the simulation include *g*\*2 alleles per epitope and *g*\*20 combinations of these genes in the gene pool. A typical simulation starts by initializing the local parasite population via a given number of transmission events with migrant repertoires whose composition is sampled randomly from a regional pool of genes, *G*. Specifically, 20 hosts are infected with randomly assembled repertoires, and one migrant repertoire is introduced every day into the population to simulate exposure to all the variants in the gene pool quickly.

### Mutation and ectopic recombination during the asexual blood stage of infection

Mutations occur at the level of epitopes. While infecting a host, each epitope can mutate at a rate *µ*_*m*_, to a new allele so that the total number of alleles, *n*, increases by one. *var* genes often change their physical location and form new variants through ectopic recombination and gene conversions. These processes occur during both sexual and asexual stages. As ectopic recombination is observed more often during the asexual stage where parasites spend most of their life cycle, and our model does not represent the mosquito stage explicitly, we consider ectopic recombination among genes within the same repertoire during the asexual stage. Two genes are selected from the repertoire, with a breakpoint located along the gene randomly. Newly recombined genes have a probability *P*_*f*_ to be functional (i.e., viable), defined by the similarity of the new variant with its parental genes as in He et al. [13]. Specifically, 

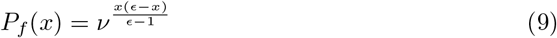

(Eq.3 in Drummond et al. [46]), where *x* is the number of mutations between the recombined gene and one of the parental genes, *E* is the genetic difference between the two parental genes and *ν* is the recombinational tolerance. If the recombined gene is selected to be non-functional, then the parental gene is kept. Otherwise, the recombined gene substitutes the parental one and a new strain is formed. In the current implementation, each recombination has a 50% chance to generate a new allele.

### *var* gene groups and trade-offs

In the model described in He et al. [13], genes differed antigenically but not functionally. For increased realism, each gene is assigned to either *var* upsGroup A or *var* upsGroup B/C to represent existing differences in recombination rates and functionality of *var* gene groups [47]. Ectopic recombination is only allowed to occur within each group, and genes in upsGroup B/C have higher recombination rates than those in upsGroup A [10]. Each gene is also assigned an intrinsic growth rate of the parasites *f* that express it (Table S2), because antigens that better bind to host receptors result in a higher parasite growth rate [48, 49]. In an additional implementation of the model, genes with higher growth rates are expressed first, followed in decreasing order by genes of lower growth rates. Also, genes with higher growth rates are expected to be cleared faster by the immune system, translating into a higher switch rate to the next gene, which is controlled by the trade-off parameter *t*_*fd*_.

## Supporting information

combined SI info

## Data Availability

Data used to produce figures are deposited in https://figshare.com/projects/An_antigenic_diversification_threshold_for_falciparum_malaria_transmission_at_high_endemicity/95803. The original C++ code for the var evolution model is available on Github (https://github.com/pascualgroup/VarModel2).

## Supporting information

### S1 Appendix. Supplementary text for theoretical derivations

**S1 Fig. Comparison between theoretical expectations from Eq. S1 (***◊***) and corresponding values from stochastic simulations (•) for** *N* **(A) and** 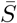 **(B) as a function of contact rate**, *β*, **repertoire size**, *g*, **and two levels of diversity ratio**, *G/g* = 10 **or 100**.

**S2 Fig. Relationship between transmission intensity and** *R*_*div*_.

**S3 Fig. The deterministic trajectory of a new gene variant invading a system that is previously at equilibrium under a low (0.015, left panels) and a high (0.05, righ panels) contact rate**.

**S4 Fig. Phase diagram of** *x*(*t*) **and** *S*_*new*_(*t*) **from S4 Fig.**

**S5 Fig. Persistence of new genes according to** 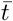.

**S1 Table. Epidemiological and genetic parameters used in stochastic simulations**.

**S2 Table. Epidemiological, genetic and within-host dynamics rules varied in the stochastic simulations**.

## Acknowledgments

We are grateful to the funding provided by the joint NIH-NSF-NIFA Ecology and Evolution of Infectious Disease (award R01 AI149779). We acknowledge valuable discussions at the Santa Fe Institute, as part of a working group supported by the James S. McDonnell Foundation (JSMF). We thank insightful comments from Karen P. Day and three anonymous reviewers on an earlier version of the manuscript. We appreciate the support of the University of Chicago through computational resources at the Midway cluster.

