## Supplementary material for "An antigenic diversification threshold for falciparum malaria transmission at high endemicity": combined SI info

### Analytical derivation of $\bar{t}$

As described in he Methods, we can derive analytically the average lifetime of a gene by assuming that the system has reached an equilibrium, including for the parasite population size $N$ and the average proportion of hosts $\bar{S}$ susceptible to an average gene. We first derive here $N$ and $\bar{S}$ based on a simplified transmission model, since these quantities will be then needed for the model of $\bar{t}$. This model is an adapted diffusion equation [1] that considers frequency-dependent selection acting upon a new gene variant entering the system at equilibrium.

#### $N$ and $\bar{S}$ at equilibrium

We first examine transmission dynamics at its equilibrium before the introduction of new genetic variants. For this we follow the dynamics of two variables: the ratio of the parasite population to the host population, $y$, and the population-level proportion of susceptible hosts to an average gene, $\bar{S}$. We consider an idealized transmission model, in which hosts keep receiving new infections at a contact rate $\beta$, up to a carrying capacity $c$, and each infection gets cleared at rate $1/\bar{\tau}$, where $\bar{\tau}$ is the average duration of infection. Concomitantly, the average proportion of hosts that are naive to a gene, $\bar{S}$ decreases as the population of hosts acquires immunity to a given gene, and increases due to the loss of specific immunity at a rate $\delta$. To calculate the rate at which a host gains immunity to a particular gene, we consider the probability that the gene is present in a parasite that is being expressed. Given that the probability that a particular gene is within a strain is $g/G$, the probability that a susceptible host is infected by a parasite genome that contains this gene is $y\bar{S}g/G$. The probability that the gene is currently being expressed is equal to the inverse of the product of the number of genes to which the host remains susceptible $((g-1)\bar{S}+1)$ and the gene switching rate $1/d$. Thus, the joint dynamics of $y$ and $\bar{S}$ can be described by the following system of differential equations

$$\begin{aligned}\frac{dy(t)}{dt} &= \bar{\beta}y(\frac{c-y}{c}) - \frac{1}{\bar{\tau}}y = \bar{\beta}y(\frac{c-y}{c}) - \frac{1}{\bar{S}gd}y \\ \frac{d\bar{S}(t)}{dt} &= -\bar{S}y\frac{g}{G}\frac{1}{(g-1)\bar{S}+1}\frac{1}{d} + \delta(1-\bar{S}) \end{aligned} \quad (1)$$

Here, $c$, the maximum number of parasites each host can sustain, is determined by either the maximum multiplicity of infection [MOI] per host (e.g., we set $max[MOI] = 10$ in the simulation), or the ratio between the diversity of the gene pool

($G$) and the genome size ($G$). Since the total infection length of a parasite is determined by its number of unique genes, the maximum number of effective parasites each host can be infected by at the same time is $c = G/g$ if each strain is composed of completely different genes, or the maximum MOI if $G/g$ exceeds this value. From this system of equations, we can derive the equilibrium values of $y^*$ and $\bar{S}^*$ (Fig. S1).

#### Derivation of $\bar{t}$ using a diffusion approximation

The expected time to fixation or loss for a newly generated variant in a population, $\bar{t}$, can be derived using a backward Kolmogorov equation following Ewens [1] Eq. 4.19, expanded to consider two dimensions for the frequency of the new variant $x(t)$ and the proportion of hosts susceptible to it, $S$,

$$
\begin{aligned}
& a(x)\frac{\partial \bar{t}(x,S)}{\partial x} + \frac{1}{2}b(x)\frac{\partial^2 \bar{t}(x,S)}{\partial x^2} + c(S)\frac{\partial \bar{t}(x,S)}{\partial S} = -1 \\
& \bar{t}(0,S) = 0
\end{aligned}
\tag{2}
$$

where $a(x)$ is the mean change of $x(t)$ in $\delta t$, $b(x)$ is the variance of $x(t)$ in $\delta t$, and $c(S)$ is the mean change of $S(t)$ in $\delta t$. As $S(t)$ directly depends on $x(t)$, it does not have a variance term.

To complete the above diffusion equation, we need to specify how $x(t)$ and $S(t)$ change in time. A new variant increases its frequency $x(t)$ at a rate that is governed by its selective advantage ($\sigma(t)$) and is scaled by its birth and death rates ($1/\bar{\tau}$). With the accumulation of population-level immune memory of the new variant as its frequency increases, its proportion of susceptible hosts decreases ($S(t)$), until it reaches the same level as that for other, older, genes ($\bar{S}$). We can describe these changes deterministically with the following dynamical system for $x(t)$ and $S(t)$,

$$
\begin{aligned}
\frac{dx(t)}{dt} &= \sigma(t)\frac{1}{\bar{\tau}}x(t)(1-x(t)) \\
\frac{dS(t)}{dt} &= -x(t)yS(t)\frac{1}{\bar{\tau}} + \delta(1-S(t)) \\
\sigma(t) &= \frac{S(t)-\bar{S}}{\bar{S}g} \\
\bar{\tau} &= dg\bar{S}
\end{aligned}
\tag{3}
$$

From numerical simulation of this system, we observe that the dynamics of $x(t)$ can be separated into two phases (Fig. S3). The first phase is a fast increasing phase resulting from its selective advantage over other genes; the second one is its stationary phase when the average proportion of susceptibles ($S$) equals that of the other genes ($\bar{S}$), and genes are maintained at a constant frequency $p = g/G$ under negative frequency-dependent (immune) selection.

We can now rewrite (2) by considering the system in (3), for the average time to absorption $\bar{t}(x,S)$ in units of $N\bar{\tau}$ as,

$$
\begin{aligned}
a(x)\frac{\partial \bar{t}(x,S)}{\partial x} + \frac{1}{2}b(x)\frac{\partial^2 \bar{t}(x,S)}{\partial x^2} + c(S)\frac{\partial \bar{t}(x,S)}{\partial S} &= -1 \\
a(x) = N\bar{\tau}\frac{dx(t)}{dt} &= N\sigma(t)x(1-x) \\
b(x) = x(1-x) & \\
c(S) = N\bar{\tau}\frac{dS(t)}{dt} &= -NxyS + N\delta(1-S)\bar{\tau} \\
\bar{t}(0, S_{new}) = 0 & \qquad (4)
\end{aligned}
$$

An explicit solution for (3) cannot be achieved because $S(t)$ changes continuously with $x(t)$. We can approximate the dynamics of the system by considering a simpler immune selection model in which $S(t)$ decreases linearly with $x(t)$ (Fig. S4), so that $\sigma(t)$ becomes negative to pull $x(t)$ back to the equilibrium frequency $x^* = p = g/G$. With this approximation, the diffusion equation in (4) then becomes,

$$
\begin{aligned}
& a(x)\frac{\partial \bar{t}(x)}{\partial x} + \frac{1}{2}b(x)\frac{\partial^2 \bar{t}(x)}{\partial x^2} = -1 \\
& a(x) = N\sigma(0)(p-x)x(1-x) = N\frac{1-\bar{S}}{\bar{S}gp}(1-\frac{x}{p})x(1-x) \\
& b(x) = x(1-x) \\
& \bar{t}(0) = 0 \qquad (5)
\end{aligned}
$$

Because genes mutate and multiple genes continuously compete for hosts, a new variant will never reach fixation and $x(t) = 0$ is its only fate. Thus, given (5), the average time for the new variant to go extinct $\bar{t}$ is given by

$$
\bar{t}(x = x_0) = 2\left[\int_0^{x_0} \frac{dy}{b(y)\Psi(y)}\int_0^y \Psi(z)dz + \int_{x_0}^1 \frac{dy}{b(y)\Psi(y)}\int_0^{x_0}\Psi(z)dz\right] \qquad (6)
$$

where $\Psi(x) = \exp\left\{\frac{N\sigma}{p}x(x-2p)\right\}$. We calculated $\bar{t}$ from both numerical integration of the above equation, and an adapted approximation based on [2],

$$
\bar{t}(\frac{1}{N}) \approx \exp\left\{N\sigma p - \frac{3}{2}log(N\sigma) - log(p) - 2\sqrt{\pi} - \frac{1}{2}log(2\pi)\right\} \qquad (7)
$$

(7) shows that $\bar{t}$ increases exponentially with the product of $N\sigma$ and $p$ when they are large (Fig. S5A). The consistently maintained stable frequency is evident in the gene trajectories of the stochastic simulations (Fig. S5B). Thus, once established, a new antigen can be maintained in a population for a much longer period than that characteristic of neutral processes.

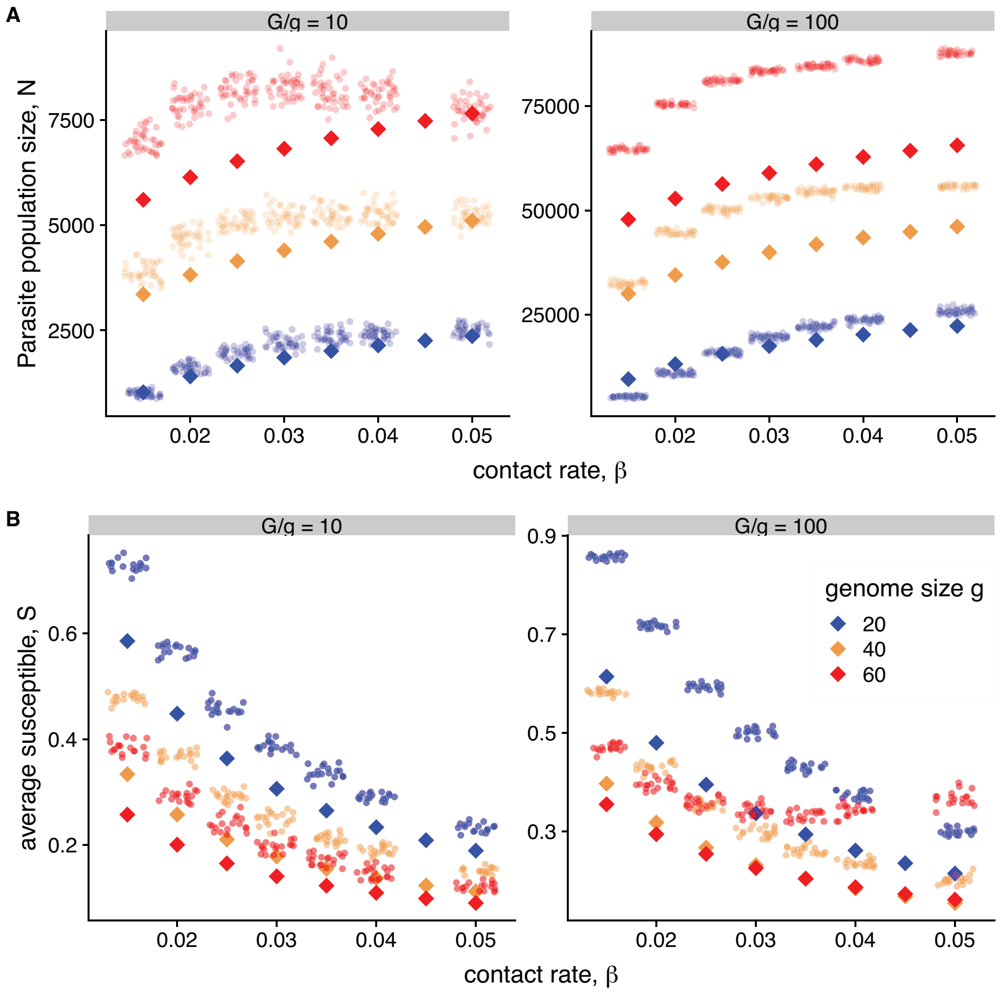


**Figure S1.** Comparison between theoretical expectations from (1) (•) and corresponding values from stochastic simulations (◇) for $N$ (**A**) and $\bar{S}$ (**B**) as a function of contact rate, $\beta$, genome size, $g$, and two levels of diversity ratio, $G/g = 10$ or 100. The simplified model predicts $N$ and $\bar{S}$ better for smaller $g$ and $G/g$, because under these conditions, the non-random arrangement of genes into parasite genomes plays a lesser role. This non-random arrangement arises from frequency-dependent selection as described in [3] which reduces overlap between among parasites. The elevated $\bar{S}$ from the simulations compared to the theoretical predictions can be attributed to the increased fitness brought by such strain structure.

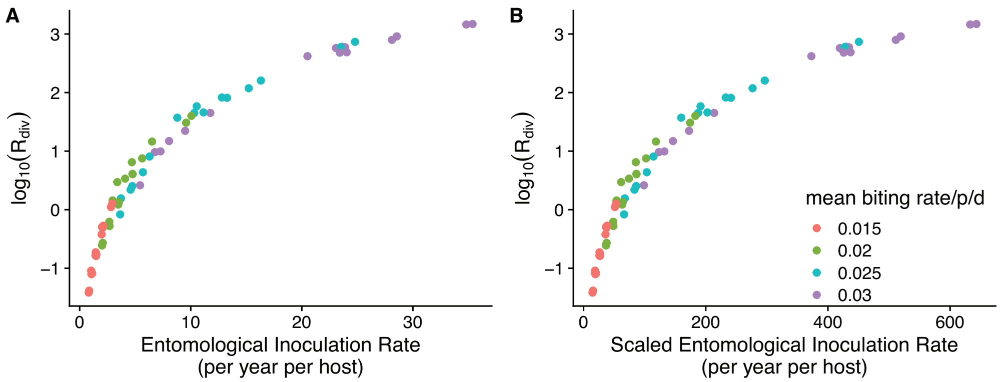


**Figure S2.** Relationship between transmission intensity and $R_{div}$. (**A**) The value of $R_{div}$ is associated with transmission intensity as measured here by the entomological inoculation rate (EIR, the number of infectious bites per person per year). (**B**) For simplicity, when a transmission event occurs, our model considers that all bites of an infected 'donor' host result in infectious bites, and that all infectious bites of the 'recipient' host result in infection. These two probabilities can be considerably less than 1 in the real world. In particular, the probability of a mosquito developing sporozoites from a blood meal ranges from 3 to 80% [4, 5], while a bite with an adequate volume of sporozoites has a probability of infecting human hosts of around 10% [6]. Thus, to compare EIR values in our model to those from the field, we must rescale them and divide them by the product of the competence/transmissibility probabilities. Here we showed an example scale that assumes the probability of mosquito developing sporozoites from a blood meal is 0.55, while a bite with an adequate volume of sporozoites has a probability of infecting human hosts of 0.1. The resulting range of EIR encompasses from low to high values, the empirical estimates from South America to Africa.

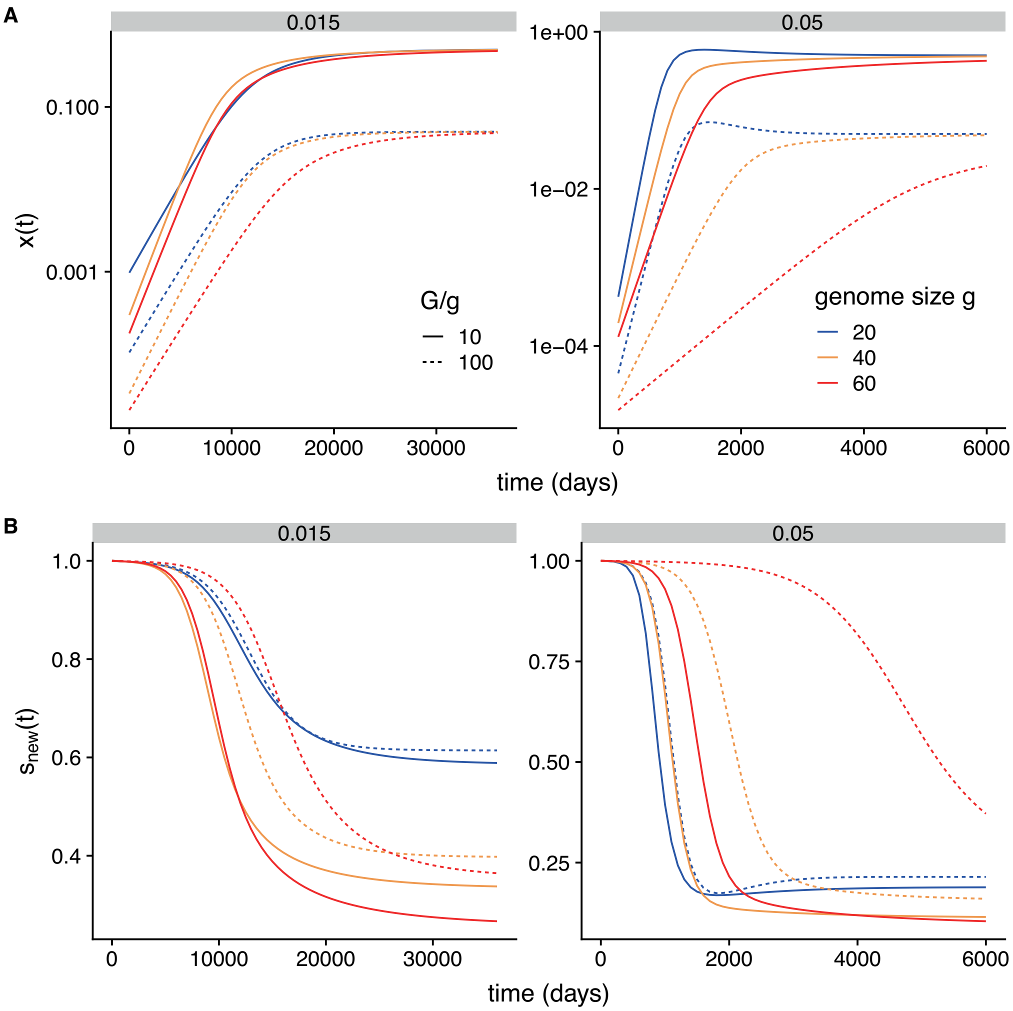


**Figure S3.** The deterministic trajectory of a new gene variant invading a system that is previously at equilibrium under a low (0.015, left panels) and a high (0.05, righ panels) contact rate. Panels in row (**A**) show the temporal dynamics of the frequency $x(t)$ , and those in row (**B**) those of which the number of susceptible hosts $S_{new}(t)$. $x(t)$ increases exponential as $S_{new}(t)$ decreases slowly in the first stage of the dynamics; it then plateaus as $S_{new}(t)$ quickly decreases.

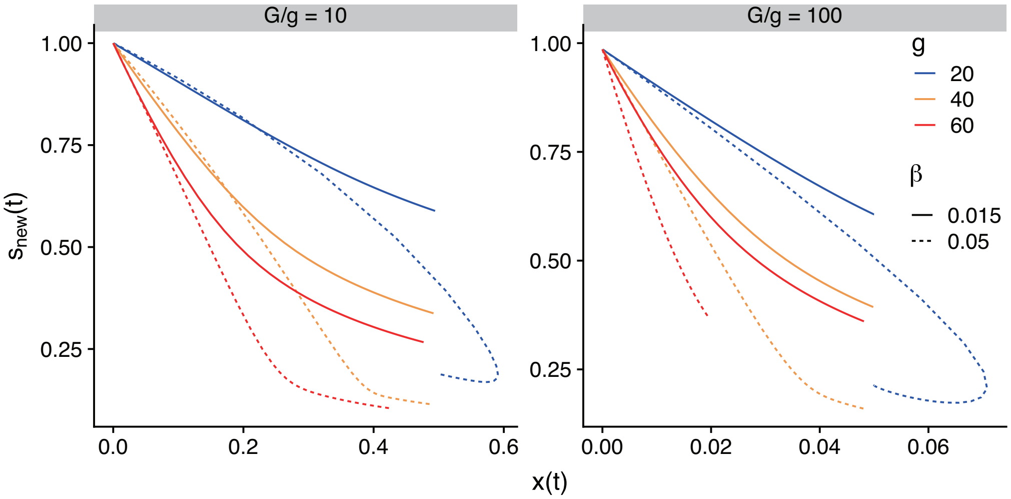


**Figure S4.** Phase diagram of $x(t)$ and $S_{new}(t)$ from Fig. S3. Because (4) does not have an explicit solution, we approximate the decrease of $S_{new}(t)$ to $\bar{S}$ as a linear function of $x(t)$. As shown here, this approximation is quite accurate when $x(t)$ is low.

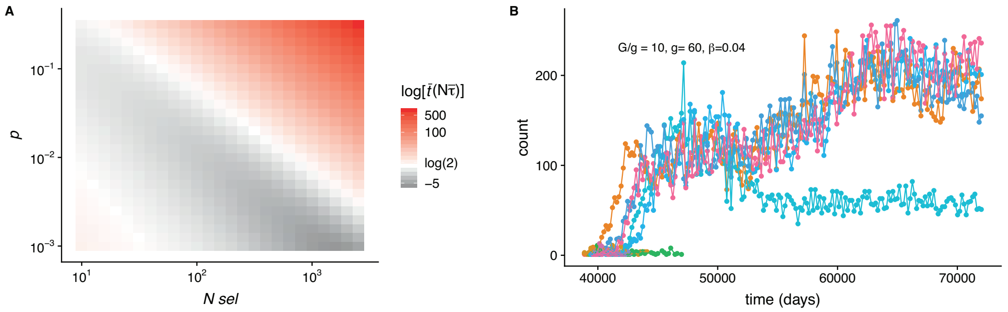


**Figure S5.** Persistence of new genes according to $\bar{t}$. As shown in (7), the average lifespan of a new antigen-encoding gene increases exponentially with the product $N\sigma p$. The color gradient in **A** represents $\bar{t}$ in units of $N\bar{\tau}$ birth-death events. White coloring indicates the parameter range where the lifespan of a new gene is equivalent to the time to fixation of a neutral gene. When new genes are highly favoured (**B**), they quickly replicate and persist at a constant frequency for a long period of time.

**Table 1.** Epidemiological and genetic parameters used in stochastic simulations.

| Symbol | Type | Description | Values | unit |
|---|---|---|---|---|
| $g$ | number | genome size | [20,40,60] | |
| $G$ | number | initial gene pool size | $g$*20 | |
| $d$ | time | length of gene expression | 6 | days |
| $\beta$ | rate | contact rate | [0.015-0.5] | per person per day |
| $\delta$ | rate | immune memory loss rate | 0.001 | per allele per day |

**Table 2.** Epidemiological, genetic and within-host dynamics rules varied in the stochastic simulations.

| Rules | Values | Description |
| --- | --- | --- |
| seasonality | non-seasonal | $\beta$ is constant |
| | seasonal | $\beta(t) = \beta\left(1 + 0.8\cos\left(2\pi\left(\left(\frac{t}{year}\right) - 0.5\right)\right)\right)$ |
| transmissibility of genes, $f$ | 1 | 100% chance of transmission to mosquito |
| | 0.5 | 50% chance of transmission to mosquito |
| trade-off between $f$ and $d$, $t_{fd}$ | [0-1] | $d_{gene} = d(f \times t_{fd} + (1 - t_{fd}))$ |
| gene expression | 1 | genes of higher functionality express first |
| | 0 | gene expression is randomly ordered |
| ratio of recombination rates | [1e-9 - 1] | ratio of ectopic recombination rates between var upsGroupA and upsGroupBC |
